# Functional Impact of CYFIP2 RNA Editing on Actin Regulation, Axon Growth, and Spinogenesis

**DOI:** 10.1101/2025.03.04.641430

**Authors:** Luca La Via, Elona Ndoj, Matteo Bertoli, Veronica Mutti, Giulia Carini, Alice Filippini, Federica Bono, Chiara Fiorentini, Giovanni Ribaudo, Alessandra Gianoncelli, Giuseppe Borsani, Isabella Russo, Alessandro Barbon

## Abstract

Cytoplasmic FMRP Interacting Protein 2 (CYFIP2) a component of the Wave Regulatory Complex (WRC), one of the most important players in regulating cellular actin dynamics. Interestingly, CYFIP2 transcript undergoes RNA editing, an epitranscriptomic modification catalysed by ADAR enzymes, that leads adenosine (A) to inosine (I) deamination. *CYFIP2* editing in the coding sequence results in a K/E substitution at amino acid 320. The functional meaning of this regulation is still unknown. In this study, we aim at investigating the potential implication of CYFIP2 RNA editing related to actin dynamics during cell differentiation, axon development and synaptogenesis in neural cells. We have generated SH-SY5Y neuroblastoma cell lines in which *CYFIP2* gene has been functionally inactivated by CRISPR-Cas9 technology. *CYFIP2* KO cells showed profound actin filaments disorganisation and loss of the capability to differentiate into a neuronal-like phenotype. Overexpression of both CYFIP2 unedited (K) and edited (E) isoforms rescued normal capability. Finally, we took advantage of primary neuronal culture where endogenous CYFIP2 was knocked down by shRNA technology and *CYFIP2* editing variants were overexpressed. While CYFIP2 KD cells reported a decrease in axon development and spine frequency, CYFIP2-E variants increase the number of axon branches, total axon length and dendritic spine frequency compared to either CYFIP2 KD cells or CYFIP-K variants. Overall, our work reveals for the first time a functional significance of the CYFIP2 K/E RNA editing process in regulating the spreading of neuronal axons during the initial stages of in-vitro development and the process of spinogenesis.

## Introduction

A variety of eukaryotic cell characteristics, including cell shape, migration and vesicle trafficking, are driven by the dynamic rearrangement of the actin cytoskeleton [1]. Coordination of actin filament dynamics is crucial for the development, maturation, and function of neurons, one of the most remarkable examples of cellular polarisation [2]. Moreover, the remodelling of the underlying actin cytoskeleton determines how dendritic spine size and shape vary in relation to excitatory synapse functions. Spine development, plasticity and synaptic function all depend on actin dynamics [3]. In addition to synaptic impacts, regulation of actin cytoskeleton dynamics is essential for the development of brain circuits complexity, modifying and guiding the extension of axonal projections from neurons to their specific destinations for synaptic connections [4].

Cytoplasmic FMRP Interacting Proteins (CYFIPs) are part of the Wiscott-Aldrich syndrome protein family verprolin-homologous protein (WAVE)-related complex (WRC), one of the most important players that regulate cellular actin dynamics. In the basal state, WRC is inhibited by CYFIPs due to the sequestration of the activity-bearing VCA domain of WAVE. Binding of the active form of the small GTPase RAC1 to CYFIPs induces conformational changes in the WRC, which releases the VCA domain to trigger ARP2/3-induced actin polymerization [5]. Accordingly, the WRC signalling pathway plays a crucial role in important neurodevelopmental processes such as axon guidance and synapse morphology.

The two family members CYFIP1 and CYFIP2, beyond being part of the WRC, might interact with the Fragile X Messenger Ribonucleoprotein (FMRP), an RNA-binding protein with an important role in translational control, the lack of which leads to fragile X syndrome (FXS) [6]. These two proteins are highly similar and conserved in several organisms [7] and share 99% identity with their orthologues in mice [6]. Of the two, CYFIP1 has been extensively studied and plays an important role in neurodevelopment by linking FMRP-dependent local translation with signalling-dependent actin remodelling through the WRC [8]. Due to their high sequence similarity, it has been proposed that CYFIP1 and CYFIP2 may share common functions. However, CYFIP1 is expressed in most tissues while CYFIP2 is highly expressed in the brain; furthermore, in the brain CYFIP1 is expressed by non-neuronal cells while CYFIP2 is more enriched in neurons than in other cell types [9]. Recently, de novo CYFIP2 mutations have been identified in patients with neurodevelopmental disorders [9], [10,11]. Finally, CYFIP2 function can be regulated at the co-transcriptional level by RNA editing, a mechanism mediated by adenosine deaminases acting on RNA (ADARs) [13]. ADAR enzymes are responsible for the modification of adenosine (A) into inosine (I). Considering that I can be read as G by the ribosome, A-to-I editing can change the amino acid insertion during translation. RNA editing in CYFIP2 results in a K320E substitution which mainly occurs in the brain. At the protein level, the editing-dependent amino acid change is located between the WAVE1 interaction domain and the FMRP interaction domain, and this modification is conserved across different species, highlighting a functional relevance that has never been investigated [14].

Here, we aim to investigate the biological implications of *CYFIP2* RNA editing related to actin dynamics during axon development and synaptogenesis in neural cells. Understanding the role of CYFIP2 RNA editing substitution would be fundamental for further analysis in both normal and pathological brains.

## Material and Methods

### SH-SY5Y culture and differentiation

The human neuroblastoma SH-SY5Y cell line was purchased from the American Tissue Culture Collection (ATCC™, CRL2266). Cell lines were grown in 1:1 mixture of Ham’s F12 Nutrient Mix (Thermo Fisher Scientific) and Dulbecco’s modified Eagle’s medium, (DMEM, Merck) containing high glucose (4.5 g/L), 4 mM L-glutamine (Thermo Fisher Scientific) and 1 mM sodium pyruvate (Thermo Fisher Scientific). The medium was supplemented with 10% v/v heat-inactivated foetal bovine serum (FBS, Thermo Fisher Scientific) and 1% Penicillin-Streptomycin solution (P/S, Merck). Cells were cultivated at 37 °C with 5% CO2 at saturated humidity and kept below ATCC passage + 15 to avoid cell senescence. To differentiate SH-SY5Y cells, a two-step retinoic acid (RA, Thermo Fisher Scientific) and brain-derived neurotrophic factor (BDNF, Merck) protocol was used [15]. In brief, cells were seeded at a density of 1 × 10^4^ cells/cm^2^ on Poly-D-Lysine (Merck) pre-coated slides. Cells were grown in RALS medium (DMEM/F12 1:1; 3% FBS; P/S; 4 mM L-glutamine; RA 10 μM). After 6 days, the culture medium was replaced with RANBB medium (NeuroBasal medium; B27; (Thermo Fisher Scientific) P/S; 4 mM L-glutamine; 10 μM RA; 50 ng/m BDNF) for another six days of culture. The culture medium was changed every two days.

### Primary hippocampal neuronal culture maturation and drugs treatments

All experiments complied with guidelines for the use of experimental animals issued by the European Community Council Directive 86/609/EEC and were approved by the Italian Ministry of Health (ID: 480/2017). Primary hippocampal neurons were prepared as described in Mingardi et al., 2021 [16]. Briefly, hippocampi from embryonic day 16.5 were dissociated mechanically and neurons were resuspended in Neurobasal™ medium (Thermo Fisher Scientific) supplemented with B27 (Thermo Fisher Scientific) containing 30 U/ml Penicillin, 30 mg/mL Streptomycin, 0.75 mM Glutamax (Thermo Fisher Scientific) and 0.75 mM L-Glutamine. Neurons were seeded at the density of 40.000 cells/cm^2^ on 0.1 mg/mL pre-coated Poly-D-Lysine glass coverslip or 25000 cells/cm^2^ in 0.02 mg/ml Poly-D-Lysine-coated 6-multiwell plates and maintained at 37°C under a 5% CO2 humid atmosphere. Three days after seeding, half of the medium was replaced with 24 hours-astrocyte-conditioned medium. Further, half of the medium was changed every seven days up to a maximum of four weeks. Various long-term pharmacological treatments were applied to the hippocampal neuronal cells growth in a 6-multiwell plates at DIV 14: 25mM KCl (Merck); 50 μM Glutamate (Merck) [17]. 100 μM APV (Merck) in combination with 1 μM TTX (Merck) [18]; cLTP (50 μM Forskolin (Merck), 100 μM Picrotoxin (Merck), 100 nM Rolipram (Merck), 50 ng/ml BDNF) [19]; or cLTD (25 μM DHPG (Merck)) (10.1038/nn746). After 24 hr of treatment, cells were harvested and total RNA was extracted to carry out an RNA editing level evaluation for *CYFIP2* transcript. The cells that grew on glass slides were used for live imaging microscopy at DIV4 or fixed for 20 min with a 4% Paraformaldehyde (Merck) solution in PBS, to be used in fluorescence microscopy.

### Molecular modeling

The structures for WAVE complexes, in the active and inactive form, were retrieved from the Protein Data Bank (PDB, rcsb.org, accessed on Jan 15, 2025). In particular, the models of cryo-EM structure of WAVE regulatory complex with Rac1 bound on both A and D site at the resolution of 3.00 Å (active, PDB ID 7USE) and of the cryo-EM structure of WAVE regulatory complex at the resolution of 3.00 Å (inactive, PDB ID 7USC) were used [20]. CYFIP1 was manually removed from such structures using UCSF Chimera 1.17.1 [21].

The model of CYFIP2 was generated using AlphaFold (AF-Q96F07-F1-v4, UniProt Q96F07) [21,22]. The K320E variant was produced with UCSF Chimera 1.17.1 using the *swapaa* command, which includes the Rotamers tool that allows the optimization of side chain orientation [21]. The Dock Prep tool was then used to prepare all the proteins before docking, and in particular to fix missing side chains and to add hydrogens and charges under the AMBER ff14SB forcefield. The protein-protein docking experiments were performed using the HDOCK server (hdock.phys.hust.edu.cn, accessed on Jan 15, 2025) [24], which is based on a hybrid algorithm of template-based modelling and *ab initio* free docking. According to the docking algorithm, the protein defined as the “ligand” is rotated and translated and the top 10 translations for each rotation are optimized by the iterative knowledge-based scoring function. The binding modes to the protein defined as the “receptor” are then clustered with a root mean square deviation (RMSD) cutoff of 5 Å. Modified WAVE complexes, in which CYFIP1 was removed, and CYFIP2 variants were uploaded to the HDOCK server as receptor and ligands, respectively, for the docking experiments performed using default settings. After the calculations, the docked structures and the result files were retrieved and processed using UCSF Chimera 1.17.1, which was also used to produce the artworks [21]. The scores obtained from docking studies were expressed in -kcal/mol.

### Total RNA Isolation, retro-transcription and sequencing

Neuronal cultures at different DIV or cell cultures (A-172; HeLa; HEK293T; SH-SY5Y) were washed with RNase-Free PBS and mechanically harvested in 1 mL of TRIzol™ (Thermo Fisher Scientific) and total RNA were extracted with Direct-zol™ RNA miniprep kit (Zymo Research) following manufacturer instructions. The concentration of the eluted RNA was measured on a Nanodrop™ ND-1000 Spectrophotometer (Thermo Fisher Scientific). The poly-A^+^ mRNA samples from different brain areas (Cc-corpus callosum; Hi-hippocampus; Nc-caudate nucleus; Sc-spinal cord; thalamus; Cx-cerebral cortex) derived from the brain pools of 19 normal male/female Caucasians (Clontech). 1 µg of total RNA from cell cultures or 50 ng of poly-A^+^ mRNA from human brain tissues was reverse transcribed using Moloney Murine Leukaemia Virus Reverse Transcriptase (M-MLV RT) (Thermo Fisher Scientific). To perform PCR reactions of the target genes, normalised amounts of the RT products was mixed with 12.5 μl of 2X DreamTaq™ Green PCR Master Mix (Thermo Fisher Scientific) and 1 μl of each forward and reverse primer (25 pmol/μl), to a final volume of 25 μl. PCR reaction was performed using standard thermal cycling conditions outlined below. The PCR products were further processed for Sanger sequencing.

The editing levels for *CYFIP2* mRNA were quantified by sequence analysis as previously described [25] using Discovery Studio (DS) Gene 1.5 (Accelrys Inc., San Diego, CA, USA). Primers used to amplify the cDNA sequence subjected to the RNA editing process are listed in Table 1.

### Cloning of human *CYFIP2* variants and lentiviral particles production

pUC19 expressing vector containing the coding sequence of human *CYFIP2* (HG16749-U) were acquired from Sino biological Inc. and amplified with primers containing AgeI and SalI restriction sites, used to insert the amplicon into pRRLSIN.cPPT.PGK-GFP.WPRE Lentiviral vector (#12252 Addgene). Reverse primers also contained the sequence coding HA-Tag (Table 1).

To generate the edited CYFIP2 isoform (CYFIP2 E), we perform *in vitro* mutagenesis using Phusion™ Site-Directed Mutagenesis Kit (Thermo Fisher Scientific) and primers bearing the desired nucleotide variation (hCYFIP2 c.958 A/G), in a PCR protocol that amplifies the entire plasmid template. The parental template is removed using a methylation-dependent endonuclease DpnI (Thermo Fisher Scientific), and STBL3 bacteria are transformed with the nuclease-resistant nicked plasmid. Primers used for mutagenesis and to sequence the full inserts are displayed in Table 1.

Lentiviral particles were produced as follows: HEK293T cells were plated at low passages (no more than P8) 24h before transfection at a density of 9.5×10^6^ in 150 mm dish; the medium was changed 2h before transfection. Cells are co-transfected using calcium-phosphate–DNA coprecipitation method. The plasmids mix used is composed of 3rd-generation lentiviral plasmids, containing: 7 µg of VSV-G envelope gene in pMD2.G backbone vector (Addgene #12259), 16.25 µg of packaging plasmids pCMV ΔR8.74 II Gen.Pack (Addgene #22036), 6.25 µg of pRSV-rev plasmid (Addgene #12253) and 32 µg of transfer Vector (pRRLSIN.cPPT.PGK-hCYFIP2-HA.WPRE K or E (from now on pRRL-hCYFIP2-K or E) to induced the expression of CYFIP2 K or E variants or pTweenLenti-H1-shCyfip2, to downregulate endogenous CYFIP2 protein).

The plasmid solution is made up with a final volume of 1225 µl with 0.1× Tris-EDTA (TE 0.1×)/dH2O (2:1); finally, 125 µl of 2.5M CaCl2 are added to the suspension; the mixture is maintained 5 min at RT. The precipitate is formed by adding dropwise 1250 µl of 2× HBS solution to the mixture. The suspension was added immediately to the cells following the addition of 2× HBS. The calcium-phosphate plasmid DNA mixture was allowed to stay on the cells for 14-16h, after which the medium was replaced with fresh one. After 24 h and 48 h from medium replacement the cells supernatant was collected. Then, after the ultracentrifugation at 26000 rpm/2 h at 4°C, the viral pellet was resuspended in PBS 1X.

### Protein extraction, quantification and Western Blot

Cells harvested from primary hippocampal cultures were solubilized with modified RIPA (50 mM Tris-HCl, pH 7.4, 150 mM NaCl, 1 mM EDTA, 0.25% Na-DHA, 0.1% SDS, 1% NP-40 and Roche protease inhibitor tablets) and then sonicated. A portion of the lysate was used for the bicinchoninic acid (BCA) protein concentration assay (Merck). Equal amounts of protein were applied to precast SDS polyacrylamide gels (4–12% NuPAGE Bis-Tris gels; Invitrogen) and the proteins were electrophoretically transferred to a nitrocellulose transfer membrane (GE HealthCare) for 2 h. The membranes were blocked for 60 min with 3% non-fat dry milk in TBS-T (Tris-buffered saline with 0.1% Tween-20, Merck) and then incubated overnight at 4°C in the blocking solution with the rabbit polyclonal anti-CYFIP2 (1:1000; #ab95969 Abcam), mouse monoclonal anti-GAPDH (1:10000, #MAB374 Millipore) primary antibodies, rabbit polyclonal anti-HA (1:1000 H6908 Merck). For detection, after 3 washes in TBS-T, the membranes were incubated for 1h at room temperature with IR-Dye secondary antibodies (1:2000 in TBS-T, LI-COR Biotech). Signals were detected using an Odyssey infrared imaging system (LI-COR Biotech) and quantified using Odyssey version 1.1 (LI-COR Biotech). Data are presented as the ratio of the intensity band of the investigated protein to that of the GAPDH band and are expressed as a percentage of the controls. Each condition was carried out and analysed in 3 independent primary culture dishes.

### Indirect immunofluorescence assay

Cells were fixed with paraformaldehyde 4% (Merck), washed three times with phosphate buffer saline (PBS) and permeabilized with 0.3% Triton-X-100 for 10 min at room temperature. After 1 h of saturation with blocking solution (2% BSA, 0.1% Triton-X-100, 2% FBS), cells were incubated for 1 h at RT with primary antibody (rabbit polyclonal anti-CYFIP1, 1:100, #ab154045, Abcam; rabbit polyclonal anti-CYFIP2, 1:100, #ab95969, Abcam; rabbit anti-HA, 1:150, #H6908, Merck). After three washes with PBS, the cells were incubated with the secondary antibody (Goat anti-Rabbit Alexa Fluor™ 594, #A-11012; Goat anti-Rabbit Alexa Fluor™ 488, #A-11008 - Thermo Fisher Scientific) 1:500 for 1 h at RT. The cells were washed three times with PBS and then the DAPI (Thermo Fisher Scientific) staining was performed. The coverslips were then mounted with SlowFade Gold reagent (Thermo Fisher Scientific) and observed under the fluorescent microscope.

### Generation of SH-SY5Y CYFIP2-Knock-out cell line

We used the CRISPR/Cas genome editing system to establish an SH-SY5Y knock-out (*KO)* cell line that is deficient in both *CYFIP2* alleles (*CYFIP2*^−/−^). The SH-SY5Y *CYFIP2 KO* cell line was generated by transfecting WT cells with the plasmid PX459 (pSpCas9(BB)-2A-Puro (PX459) V2.0; Addgene #62988) [26]. The plasmid contains two expression cassettes: a human codon-optimised SpCas9 and the single guide RNA which consists of 76 nt of a scaffold sequence required for Cas-binding and 20 nt of a sequence that identifies the locus in the genome to be targeted. The expression of sgRNA is under the control of the U6 promoter. After digesting the vector with BbsI (Thermo Fisher Scientific #ER1011), two annealed complementary oligos were cloned upstream the sgRNA scaffold. The oligos were selected using the CRISPOR tool Version 5.01 (http://crispor.tefor.net/) [27], based on the 20 bp target sequence and were followed on the 3’ end by a 3 bp NGG PAM sequence. We have chosen a predicted guide that has a high specificity score [28] and a low number of predicted mismatches. After cloning, Stbl3™ Chemically Competent *E. coli* strain (Thermo Fisher Scientific #C737303) was transformed with the purify PX459-CYFIP2-*KO* DNA plasmid.

SH-SY5Y cell line (ATCC™, CRL2266) was transfected with PX459-*CYFIP2*-*KO* DNA plasmid, using Lipofectamine™ 3000 (#L3000001 Thermo Fisher Scientific), following the manufacturer’s protocol. We selected transfected cells using puromycin at the concentration of 2 ng/μl for 24 hours. After DNA extraction, the *CYFIP2* gene target region for Cas9 was PCR amplified and sequenced to evaluate the presence of genome editing events. The resultant electropherograms were assessed using the ICE Analysis tool (Syntego^©^ https://ice.synthego.com/#/).

### Generation of SH-SY5Y CYFIP2 K/E Knock-In cell lines

To generate a stable population of CYFIP2 K and E (SH-SY5Y knock-in (KI cell lines), SH-SY5Y CYFIP2-*KO* cells were transduced with a MOI 1 of lentiviral particles generated with transgenic vectors pRRL-hCYFIP2-K or E as previously described, directly into cell medium. We selected transduced cells using puromycin at the concentration of 2 ng/μl for 24 hours.

### Generation of lentiviral vector to downregulate endogenous CYFIP2 protein

shRNA expressing vectors to downregulate endogenous CYFIP2 protein in hippocampal cells, were designed starting from Tween-Lenti vector (7970 bp), that also contains the Green Fluorescent Protein (GFP) coding sequence under the control of the hPGK promoter, kindly provided by Prof. Leonardo Elia (University of Brescia). Expression of shRNA sequence is under the control of H1 promoter. After the vector was double-digested with XbaI and XhoI (#ER0681; #ER0692 Thermo Fisher Scientific), two annealed complementary oligonucleotides carrying the palindromic sequence of shRNA to *Cyfip2* mRNA were cloned. The sequence of shRNA that recognise the 3’UTR sequence of mouse *Cyfip2* transcript starting from position 3775 was selected using shRNA Optimal Design tool from Kay Lab (https://med.stanford.edu/kaylab). After cloning, Stbl3™ Chemically Competent *E. coli* strain (#C737303 Thermo Fisher Scientific) was transformed with the pTweenLenti-H1-shCyfip2 DNA, used to generate Lentiviral particles as previously described.

To generate a CYFIP2 K/E KI model, hippocampal cultures at DIV1 were transduced adding a mix of lentiviral particles containing TweenLentiH1-shCyfip2 to downregulate the endogenous CYFIP2 protein and pRRL-hCYFIP2-K or E to induce the expression of CYFIP2 K or E variants. 4% PFA was used to fix neuronal cultures at DIV4 or DIV17 for morphological analysis.

### Confocal microscopy and Imaging Analysis

Fluorescently labelled cells were acquired using an inverted laser scanning confocal microscope LSM900 Axio Observer.Z1/7 (Carl Zeiss, Jena, Germany) with an EC Plan-Neofluar 40x/1.30 Oil DIC objective, 1 slice, 625 Tiles (23033x2333 pixels), Pixel Time 2.06 µs, were used for images acquisition of SH-SY5Y cells and for hippocampal cells at DIV3. For spine analysis, a Plan-Apochromat 63x/1.40 Oil objective, 19 slices (6.3µm), Pixel Time 3.55 µs were used for images acquisition of hippocampal cells at DIV18. Pictures represent a maximum intensity projection (MIP) of 19 Z sections at 0.33 µm of interval). SH-SY5H total neurite length, neuronal dendritic length and number of branches were measured using Simple Neurite Tracer from Fiji [29]. For each condition, a minimum of 15 cells were analysed. The number of spines was measured manually using ImageJ and spine density was calculated by quantifying the number of spines in a 10 µm dendritic segment. For each condition, spines of three secondary dendrites from a minimum of 15 cells were analysed.

### Statistical analysis

Data were shown as mean ± standard error of mean (SEM). Statistical analysis of the data was performed using GraphPad Prism 8 (GraphPad Software Inc., USA). For all the experiments, one-way ANOVA followed by Tukey’s post-hoc test was used. Statistical significance was assumed at p<0.05.

## Results

### 1. CYFIP2 RNA editing is conserved among organism, modulated during neuronal maturation and by neuronal activity

To get new insights in the functional role of *CYFIP1/2* in neuronal development, we have examined their protein expression in mouse primary hippocampal neuronal cultures at different Day In Vitro (DIV) as a model of neuronal maturation. CYFIP1 is barely expressed at DIV4 while it increased through the differentiation process with a peak of expression at DIV18; somehow dissimilarly, CYFIP2 is already highly expressed at DIV11 and stable till DIV18 (Fig. 1a).

**Figure 1:**
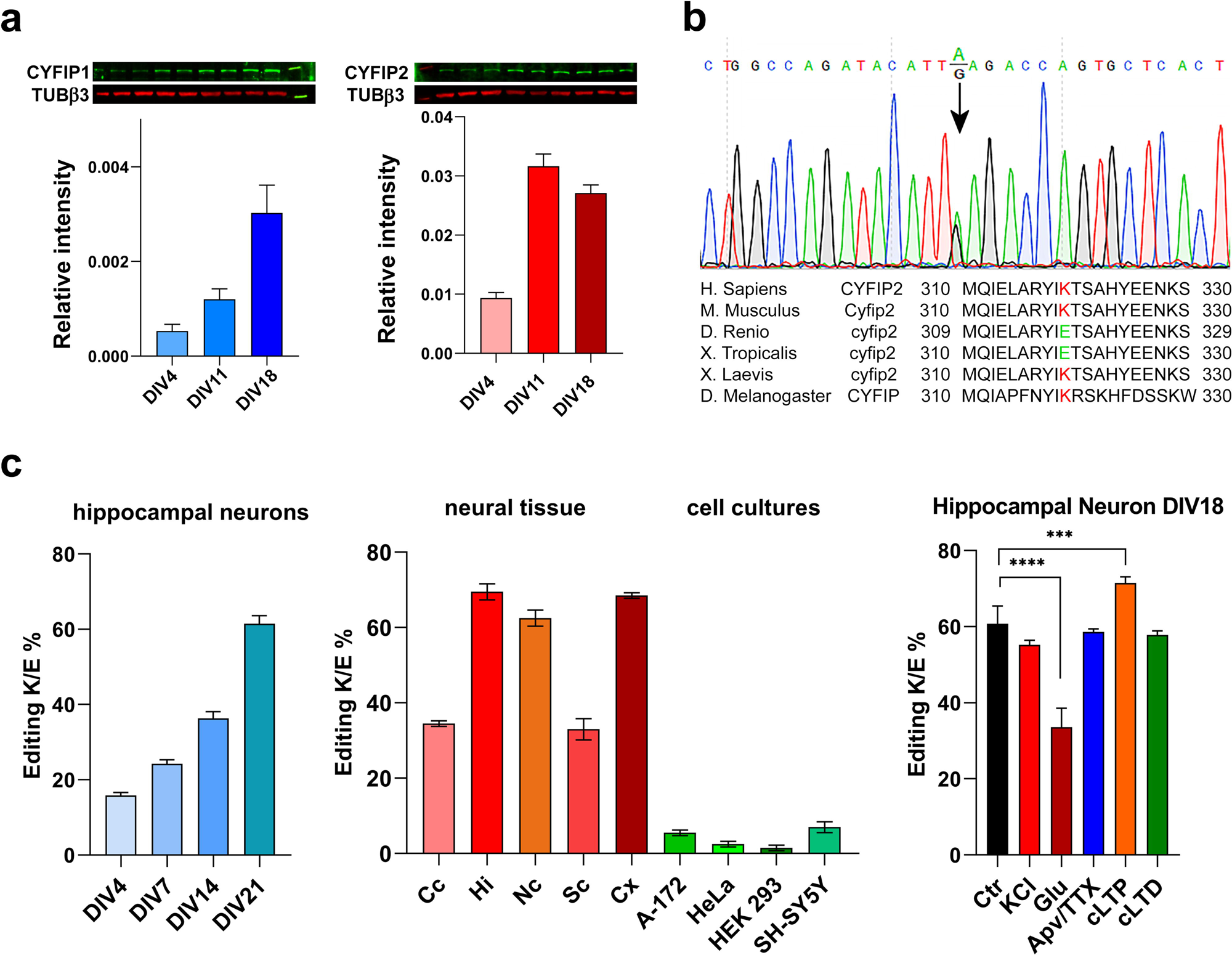
(a) CYFIP1 and CYFIP2 protein expression during neuronal maturation. (b) Electropherograms from Sanger sequencing of CYFIP2 exon 10 carrying the A/I RNA editing sites. The editing site is represented by the overlapping of A and G peaks. Alignment of human CYFIP2 protein (NP_001032410.1) against the mouse (NP_598530.2), zebrafish (NP_001091056.1), Xenopus (NP_001103523.1), Drosophila (NP_650447.1) homologs showing the conservation of sequences where the edited amino acids are located. (c) Analysis of editing levels of the CYFIP2 K/E site during *in vitro* primary hippocampal neuron maturation, in several human brain regions and cell lines, and in primary hippocampal neurons after neuronal activity modulation.

We then focused on CYFIP2 K/E RNA editing and we aligned the protein sequence of different organisms (Fig. 1b), showing the high conservation of amino acid residues where the editing event takes place. Interestingly, *Danio rerio* and *Xenopus tropicalis* encode a CYFIP2 isoform with the E variant without the intervention of RNA editing. Then we analysed RNA editing level during neuronal differentiation. Our findings demonstrated that K/E RNA editing level increased during in vitro neural development (DIV4: 15.8%; DIV7: 24.3% DIV14: 36.3%; DIV21: 61.5%) (Fig. 1c, left), indicating that the process is tightly regulated during maturation. Moreover, RNA editing levels in different human post-mortem brain regions and non-neural cell line samples have been examined (Fig. 1c, middle). The editing percentage obtained from neural tissue samples are: Corpus Callosum (CC) 35%; Hippocampus (Hi) 71%; Caudate (Nc) 61%; Spinal Cord (Sc) 31% and Cerebral Cortex (Cx) 68%. Editing percentage obtained from non-neural cell lines are as follows: glioblastoma (A-172) 6%; Human cervix epithelioid carcinoma (HeLa) 2%; human embryonic kidney (HEK293) 1%; human neuroblastoma (SH-SY5Y) 6%. Our data demonstrate that *CYFIP2* RNA editing process is specifically controlled in various brain regions and is mainly active in the central nervous system (see also, Supplementary Figure 1).

To evaluate the effects of neuronal activity on *CYFIP2* editing process, we treated mouse primary neuronal cultures with KCl and glutamate to mimic general synaptic and glutamatergic activation and with APV/TTX to mimic synaptic blocking (Fig. 1c, right). Furthermore, chemical long-term potentiation (cLTP) and long-term depression (cLTD) were induced in neuronal cells. One way ANOVA test indicates a strong effect of the treatments on RNA editing levels (F (4, 14) = 125; p<0,001). In particular, glutamate treatments lead to a downregulation on *CYFIP2* RNA editing level (Glu −55.2%; P < 0.0001). An opposite effect was found after chemical LTP induction (cLTP +17.69% p=0.0005) while cLTD did not induce variation.

### 2. Modeling of CYFIP2 editing variants and protein-protein docking

To evaluate the binding mode of CYFIP2 variants to the proteins forming the WAVE complex from a structural point of view, computational tools were employed. Structural data are already available in the literature for CYFIP1, as the structure of its WAVE complex was experimentally determined in the active and inactive states by means of X-ray diffraction or cryo-EM [21,22]. On the other hand, insights into the interaction motif of CYFIP2, and particularly of the K and E variants, are lacking.

Thus, in our study, we initially generated the model of CYFIP2 K variant using AlphaFold [21,22], and we then produced and refined the structure of the E variant as detailed in the Material and Methods section using UCSF Chimera 1.17.1 [21]. In Figure 2a, the 3D structure of CYFIP2 K variant is reported, and the residue number 320 is highlighted in red. A detailed comparison of the residues in the K and E variants is reported in Figure 2b.

**Figure 2:**
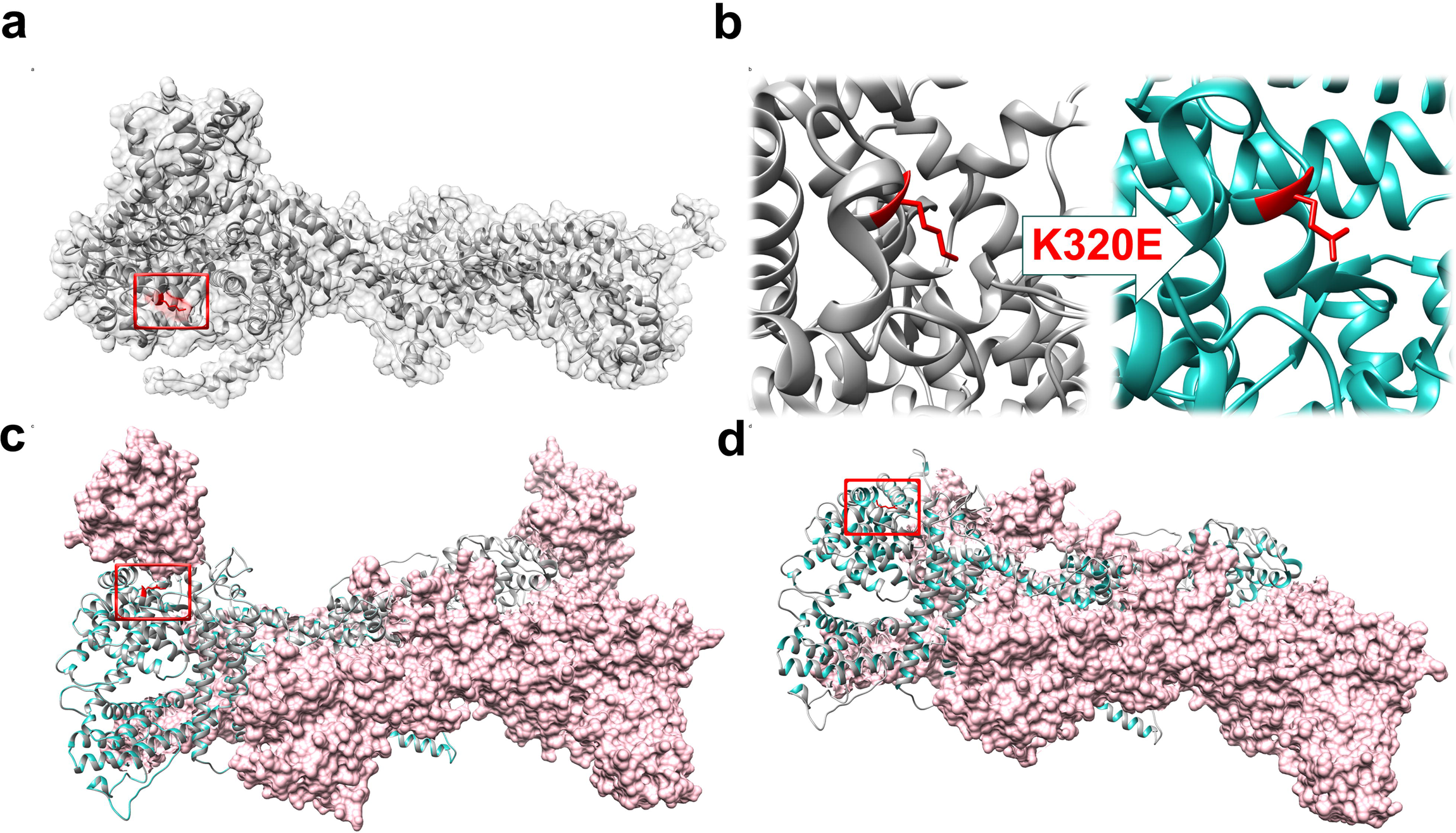
(a) 3D model of CYFIP2 built with AlphaFold; (b) detailed view of the different residues presents in the CYFIP2 K and E variants. (c) 3D model of the active and (d) inactive complexes. Residues in position 320 have been highlighted in red. The CYFIP2 K structure is depicted in grey, while the E variant in sea green. The portions depicted through pink surfaces represent the parts of the macromolecules used as receptors in the docking studies

To obtain the models of the WAVE complex with CYFIP2, protein-protein docking was used. Structural studies related to autism-linked pathological variants in the WAVE regulatory complex, based on such technique, recently appeared in the literature [30]. In our work, we selected the cryo-EM structures of WAVE regulatory complex formed by CYFIP1 as templates of the active (PDB ID 7USE) and inactive (PDB ID 7USC) forms, the latter of which lacks Rac1 [20]. The structures were edited by removing CYFIP1 from the models using UCSF Chimera 1.17.1 [21], thus obtaining the “receptor” files for docking studies. Subsequently, CYFIP2 K and E variants were used as “ligands” in the docking simulations, which were performed using the HDOCK server (hdock.phys.hust.edu.cn, accessed on Jan 15, 2025) [24]. Thus, four models of the WAVE complex were obtained through these docking runs, namely active WAVE/CYFIP2 K, active WAVE/CYFIP2 E, inactive WAVE/ CYFIP2 K, and inactive WAVE/ CYFIP2 E. These models were then ranked in terms of docking score (-kcal/mol) and inspected using UCSF Chimera 1.17.1 [21].

In the case of active WAVE complex, small differences in terms of docking scores were retrieved, as values of -621.71 kcal/mol and of -621.34 kcal/mol were computed for active WAVE/CYFIP2 K and active WAVE/CYFIP2 E, respectively. It must be pointed out that also in the case of the aforementioned study by Xie and colleagues, performed to assess the impact of pathological mutations, limited differences in terms of affinity index were detected [30]. Accordingly, from a structural point of view, it can be noted that CYFIP2 K and E variants share a similar positioning while forming the respective complexes (Figure 2c). The residue of interest is indeed located in an exposed portion of the macromolecule, pointing towards Rac1, even if the distance between this residue and the closest of Rac1 (Gln2) is of 8.4 Å.

In the case of the inactive complex, a docking score value of -1198.81 kcal/mol was computed for both inactive WAVE/CYFIP2 K and inactive WAVE/CYFIP2 E complexes. As evidenced by this result, no difference was obtained from this calculation in terms of docking score. This agrees with the model depicted in Figure 2d, where a perfect superimposition between CYFIP2 K and E variants is observed. Additionally, it must be noted that the residue of interest, in this case, is exposed to the solvent and not in proximity to other macromolecular interactions, *e.g.,* Rac1.

### 2. CYFIP2 Knock-Out induces SH-SY5Y morphological alterations that are rescued by both CYFIP2 K and E Knock-In

We decided to use a neuroblastoma SH-SY5Y cell line as an in vitro model for the study of CYFIP2 K/E editing, due to the ability of these cells to develop a neuronal phenotype. We established an SH-SY5Y KO cell line that is deficient in both *CYFIP2* alleles (*CYFIP2*^−/−^) using the CRISPR-Cas9 genome editing technology (Supplementary Figure 2) and thus does not express CYFIP2 protein (Figure 3a). Furthermore, through transduction of lentiviral particles carrying the whole CDS of human CYFIP2 K and E isoforms, KO cell line was used to create CYFIP2 Unedited (K) and Edited (E) Knock-In cell lines (Figure 3b).

**Figure 3:**
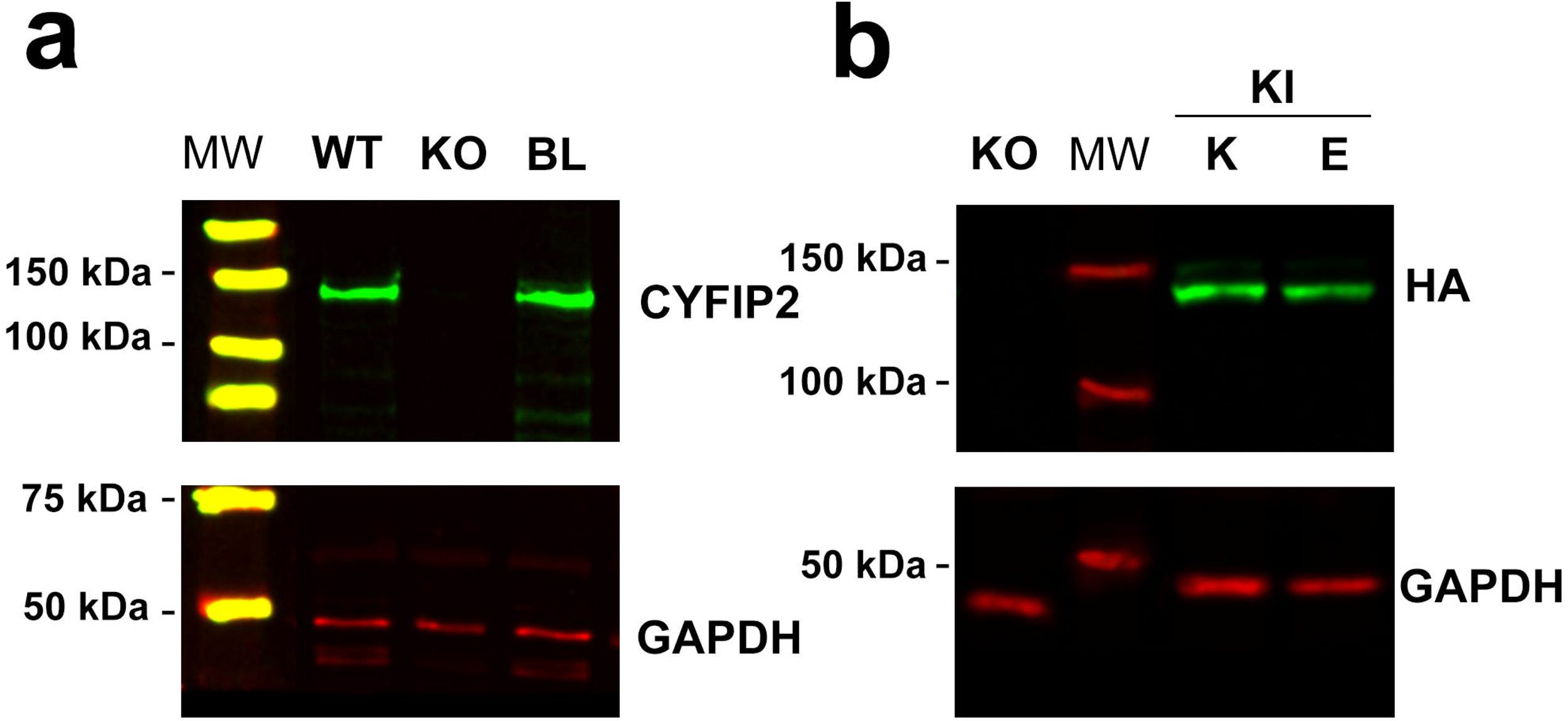
(a) IR acquisition of WB obtained from cellular lysate derived from WT SHSY-5Y cell line (WT), from SHSY-5Y cell line after CRISPR/Cas9 and puromycin selection (KO) and mouse brain lysate as a positive control. Endogenous CYFIP2 proteins are detected using rabbit-α-CYFIP2 antibody. (b) IR acquisition of WB obtained from cellular lysate derived from CYFIP2 KO SHSY-5Y cell line (KO), and SHSY-5Y CYFIP2 knock-in (KI) for the K or E variants. Exogenous CYFIP2 proteins are detected using rabbit-α-HA antibody

WT cells are characterised by a neuroblast-like phenotype with a few shortened processes (Figure 4a) while SH-SY5Y-CYFIP2 KO cells exhibit a striking morphological change with an increase in the cell area (p<0,0001 vs WT) and a decrease in the aspect ratio (major axis/minor axis p<0,0001 vs WT). Furthermore, the formation of cell blebs on the plasma membrane (Supplementary Figure 3, and Figure 3b) could be observed. In addition, the KO cells showed a completely different actin filaments organisation compared to WT cells as evidenced by the increase in stress fibers and phalloidin intensity (p<0,0001 vs WT); of note, overexpression of CYFIP2 K and E forms rescued the morphological parameters (Figure 4b) to WT levels and only differences in phalloidin intensity among the different cell lines could be observed.

**Figure 4:**
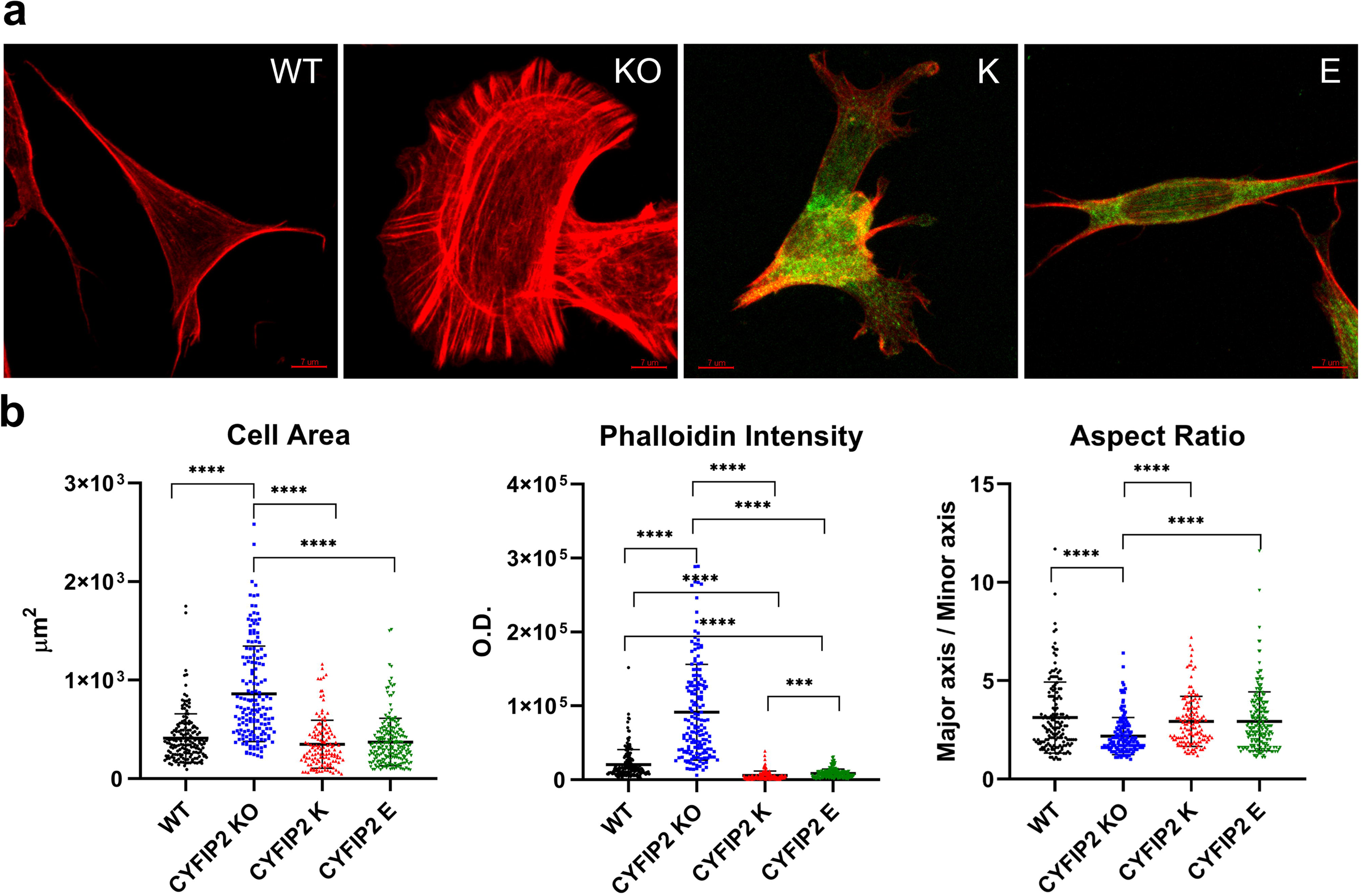
(a) CYFIP2 WT, CYFIP2 KO, overexpressing CYFIP2-K and CYFIP2-E variants SH-SY5Y cells. Actin filaments are stained by phalloidin (red) while exogenous CYFIP2 is stained with HA antibodies (green). (b) Morphological analysis of CYFIP2 WT, CYFIP2 KO, CYFIP2 K and CYFIP2 E, SH-SY5Y cell lines. The different morphological parameters (cell area, Phalloidin Intensity, Aspect Ratio) have been analysed with ImageJ. (CYFIP2 WT n = 165, CYFIP2 KO n = 161, CYFIP2 K n = 139 and CYFIP2 E n = 212. One-way ANOVA followed by Tukey’s multiple comparisons test p<0,005. Data are represented as mean ± SEM.

### 3. CYFIP2 RNA Editing modulate neurite development during SH-SY5Y differentiation

Since CYFIP2 protein is mainly expressed in neurons [9] and K/E RNA editing reaction is active only in the central nervous system [14], we investigated the role of CYFIP2 K/E variants on neuronal development. For this purpose, we used a well-established model of neuronal differentiation based on the application of a two-step retinoic acid (RA) and brain-derived neurotrophic factor (BDNF) treatment to SH-SY5Y cells (Fig. 5a) [15] to monitor the induction of undifferentiated SH-SY5Y into neuron-like cells with distinctly polarised axon-dendritic morphology. Treated cells have spindle-like morphologies with polarised appearances, with sporadic shafts and projections extending from soma, usually one per cell. When exposed to BDNF, cellular processes expanded quickly and dramatically, showing identifiable neuron-like characteristics (Fig. 5b). One-way ANOVA analysis reported statistically significant differences among the groups (F(3,795) p<0.0001) in the total neurite length (CYFIP WT mean length = 56.0 ± 3.69, n = 243; CYFIP KO mean length = 15.89+/-1.17, n = 173; CYFIP-K mean length = 53.34+/-2.89, n = 192; CYFIP-E mean length = 60.04+/-4.21, n = 191). Tukey’s multiple comparisons test showed a dramatic decrease after knocking out *CYFIP2* gene expression. Furthermore, the ability of neurite development is completely restored in both *CYFIP2* KI cell populations (K/E) (WT vs KO Mean diff. 40.14, p <0.0001; KO vs K Mean diff. -37.45, p <0.0001, KO vs E Mean diff. -44.15, p <0,0001). However, no differences could be detected between CYFIP K and E cell populations (K vs E Mean diff. -6.70, p = 0.51), indicating that CYFIP2 is important for neuronal differentiation but the two isoforms might have a similar function regarding this process in SH-SY5Y cells (Fig. 5c).

**Figure 5:**
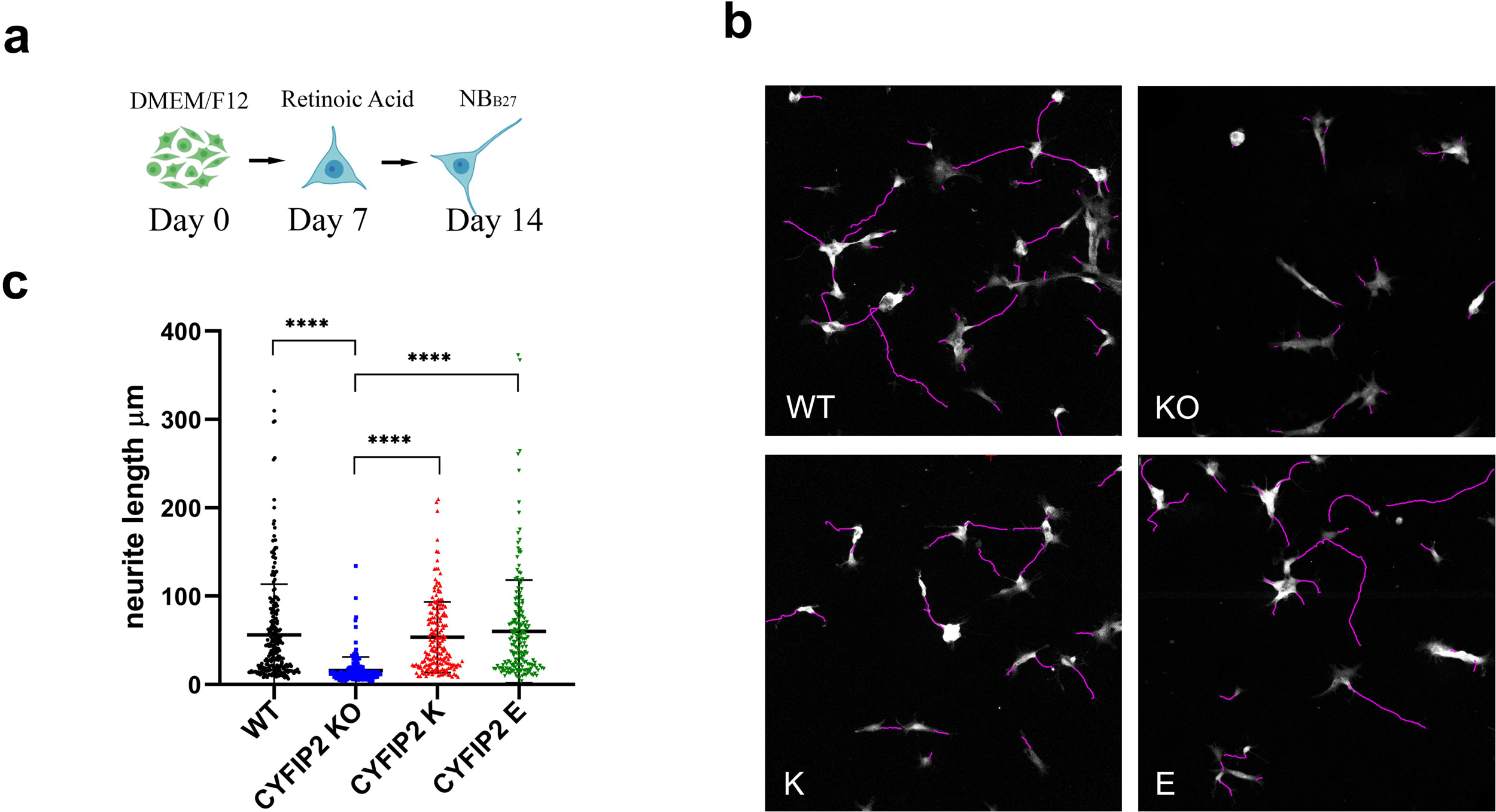
(a) Schematic representation of SHSY-5Y neuronal differentiation. (b) Representative cell images for neurite length analysis by Simple Neurite Tracing. (c) Neurite length analysis of the different populations of SHSY-5Y of cells. (One-way ANOVA followed by Tukey’s multiple comparisons test n = 243 (WT), 173 (KO), 192 (CYFIP2 K), 191 (CYFIP2 E): ****p<0.0001).

### 4. CYFIP2 RNA Editing alters axon development and spine frequency of hippocampal primary neurons

Since we did not observe a clear difference between CYFIP2 K/E variants in SH-SY5Y neuronal differentiation, we took advantage of primary neuronal culture model to understand the effect of CYFIP2 variants on neuronal maturation.

To down-regulate the expression of the endogenous CYFIP2 protein, we transduced primary hippocampal neuronal cultures at DIV1 with lentiviral particles containing a shRNA targeting the 3’UTR of the *Cyfip*2 gene. Since the vector carrying the shRNA also contains the sequence coding for green fluorescent protein (GFP), we evaluated the extent of transduction by fluorescence microscopy technique (Figure 6c). As shown in Figure 6a, 48 h after transduction, the levels of endogenous CYFIP2 protein were reduced by 70.2%. CYFIP2 knocked-down cells were co-transduced with lentiviral particles carrying plasmids expressing the coding sequence of human CYFIP2 K or E exogenous variants, to study the impact of edited or unedited CYFIP2 variants on *in vitro* neural maturation. Western blot analysis showed the similar expression of both exogenous CYFIP2 variants, eighteen days after lentiviral transduction (Figure 6b). The analysis of the two CYFIP2 KI lines showed a robust down-regulation of editing level for CYFIP2-K and a moderate upregulation for CYFIP2-E line, relative to control levels (ANOVA F(2,6) p<0.0001; WT=42.0 *±* 2.0 %; CYFIP2-K= 24.0±1.2 % p<0.001 vs Ctr; CYFIP2-E= 53.3±1.5 % p<0,05 vs CTR; p<0.001 vs K) as expected (Figure 6d).

**Figure 6:**
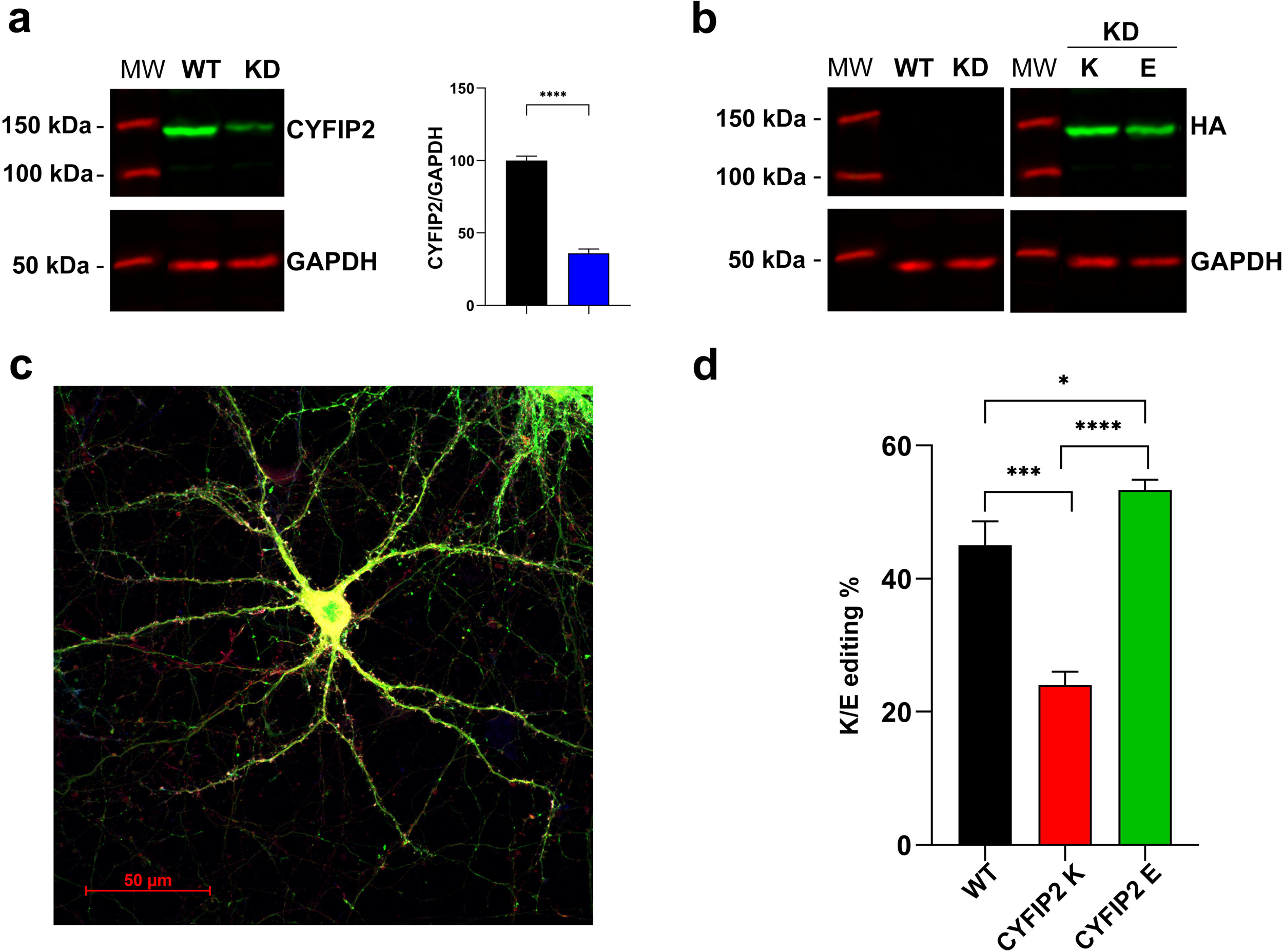
(a) IR acquisition of WB obtained from cellular lysate derived from WT Hippocampal culture (WT) and after 48 from Lentiviral transduction of shRNA targeting the 3’UTR of Cyfip2 gene (KD). Endogenous CYFIP2 protein is detected with rabbit-α-CYFIP2 antibody. In the graph CYFIP2 downregulation percentage is reported. (b) Cellular lysate of KD hippocampal cells growth for 18 DIV after lentiviral transduction with CDS of human CYFIP2 K or E variants. Exogenous CYFIP2 proteins are detected using rabbit-α-HA antibody. (c) DIV18 hippocampal cell captured by confocal microscopy. The GFP protein, which represents a marker for CYFIP2 down-regulation, is detected in green. Exogenous CYFIP2 is visible in red (rabbit-α-HA antibody -goat-α-rabbit Alexa Fluor 594). (d) CYFIP2 K/E editing levels analysed from total RNA samples obtained from wild-type hippocampal cell (WT) or CYFIP2 knock-down hippocampal cell transduced with two editing variants (CYFIP2 + CYFIP2 K; CYFIP2 + CYFIP2 E) (Tukey’s multiple comparisons test: *p<0.05 ** p<0.01 ***p<0.001 ****p<0.0001).

We initially focused on the early stages of neuronal maturation, analysing neuronal axon length and complexity during the first days of in vitro neuronal growth (Holt et al., 29518358) (Figure 7a). One-way ANOVA found a statistically significant difference among groups in each of the three parameters considered (number of branches F (3,260) p<0.0001; path length F (3,260) p< 0.0001 and complexity index F (3,260) p<0.0001). Tukey’s multiple comparisons test showed a reduction in the number of branches (WT=19.55 *±* 1.55; CYFIP2-KD 11.13 ± 0.92 p<0,001) and path length (WT=768.0 *±* 55.1; CYFIP2-KD 427.0 ± 33.3 p<0,001) after knocking down the *CYFIP2* gene (Figure 7b). Furthermore, the ability to produce axons with proper complexity is completely restored when KD neurons were grown overexpressing the E variant of the CYFIP2 gene, but not with the K variant (number of branches: CYFIP2-K: 12.45 ± 1.10; CYFIP2-E: 36.77 ± 3.49; path length: CYFIP2-K: 658.6 ± 59.4; CYFIP2-E: 1141 ± 99.6). The complexity of the axons in neurons carrying the CYFIP2 E variant is statistically significantly higher even in comparison to WT neurons (Figure 7b).

**Figure 7:**
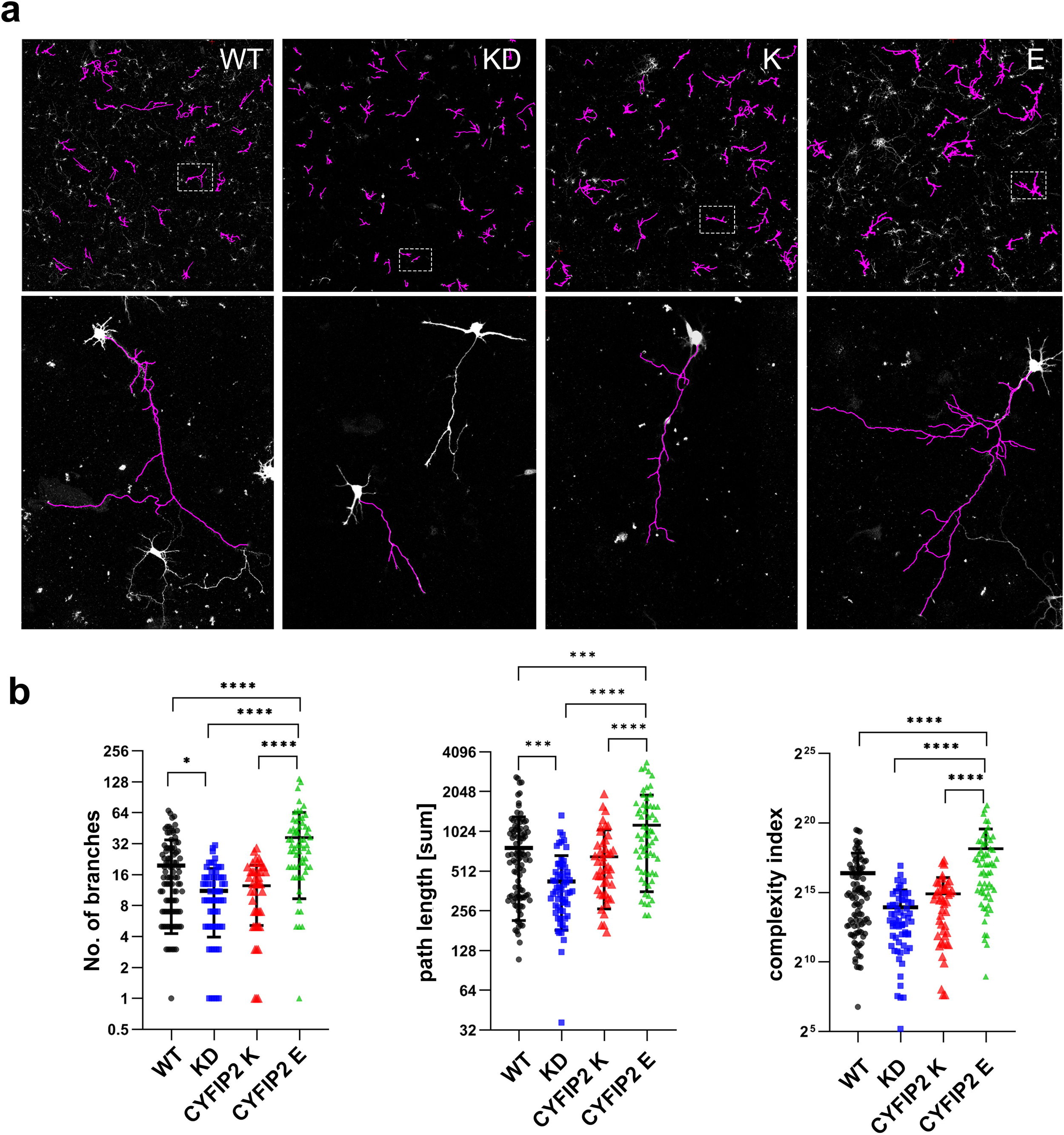
(a) Axon length analysis of hippocampal culture at DIV4 by Simple Neurite Tracing is shown in purple. (b) The complexity of axonal arborization was determined by analysing the number of branches, the total length of axons and the complexity index. (N= 97 (WT), 61 (KD), 44 (CYFIP2 K), 62 (CYFIP2 E); Tukey’s multiple comparisons test: *p<0.05; **p<0.01 ***p<0.001; ****p<0.0001).

Actin dynamics are essential for synaptic function, spine formation, and plasticity [31]. The frequency of the spines, which is the number of spines for each ten micrometres in length of the secondary dendrite (Figure 8a), significantly changed among the three different experimental groups (Figure 8b: One way ANOVA F (3,69) p<0,0001). In particular, the silencing of the CYFIP2 protein (KD), shows a decrease in the spine frequency compared to WT cells (WT: 3.65 ± 0.23; CYFIP2 KD: 2.06 ± 0.16; Tukey’s multiple comparisons test WT vs. KD p <0.0001). This decrease remains similar when the expression of the K variant was induced (CYFIP2 K: 2.50 ± 0.12; WT vs. K p<0.0001) indicating that the physiological frequency of spines is not recovered after the expression of the CYFIP2 K variant; (CYFIP2 KD vs. K ns p <0.3832). On the contrary, this parameter was completely restored after the expression of the CYFIP2 E variant (CYFIP2 E: 3.82 +/-0.26; KD vs. E p<0.0001). This difference resulted to be statistically significant also between K and E groups (CYFIP2 K vs. E p<0.0001). Taken together, the results obtained suggest a clear role of CYFIP2 K/E RNA editing process in regulating the spinogenesis process in the hippocampal neurons in vitro.

**Figure 8:**
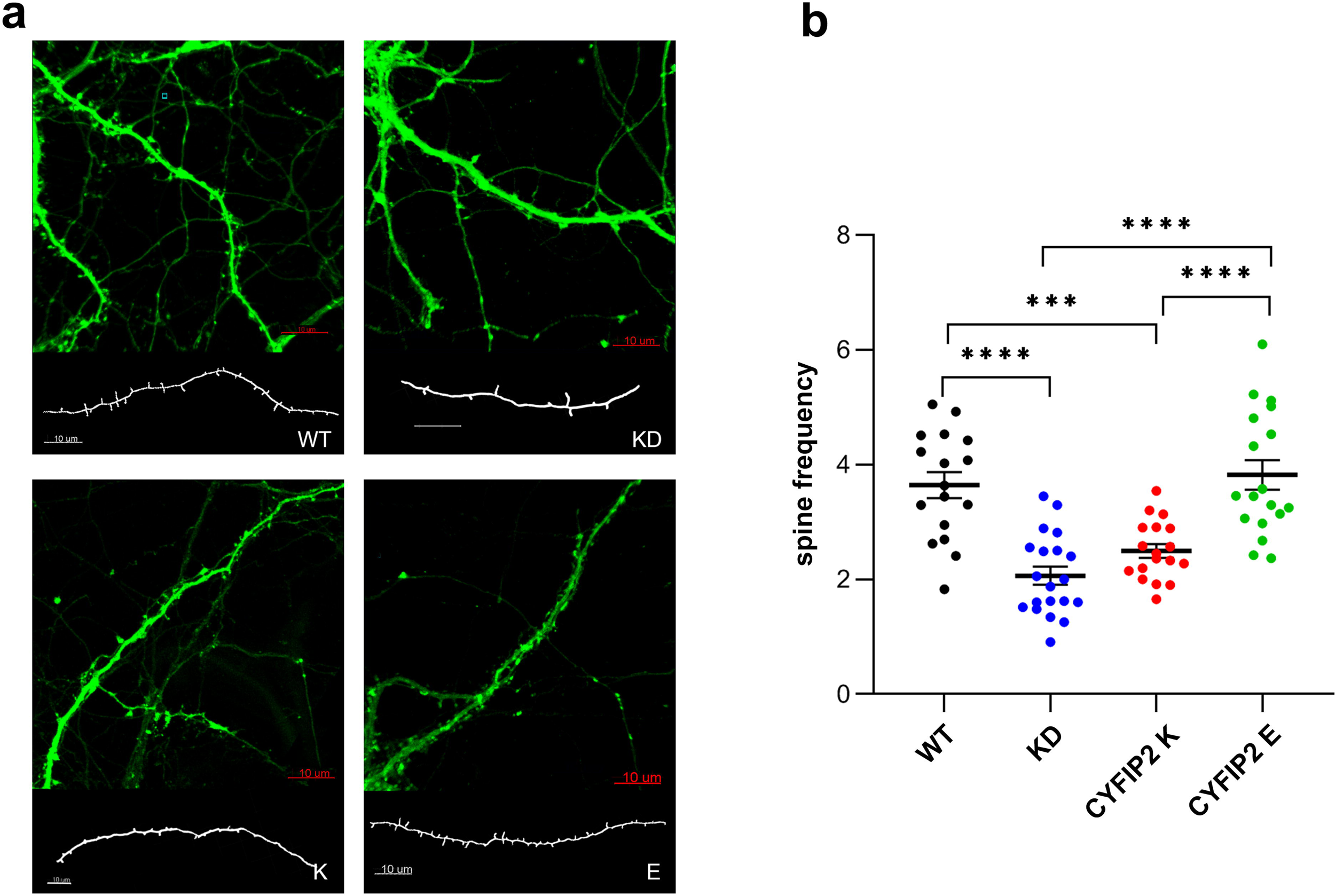
(a) Image of secondary dendrite hippocampal neurons at DIV18, captured by confocal microscopy. GFP protein signal visible in green, represents a marker of CYFIP2 down-regulation. In white, the reconstruction of dendrites and spine structures by imageJ is reported. (b) Distribution of spine frequency of the different populations of CYFIP2 cells (right) (N= 17 (WT), 20 (KD), 18 (CYFIP2 K), 18 (CYFIP2 E); (Tukey’s multiple comparisons test: ***<0.001 ****<0.0001).

## Discussion

*CYFIP2* is one of the few neuro-specific transcripts that undergo recoding RNA editing [14] resulting in a K/E substitution at amino acid 320; nevertheless, its functional meaning is still unknown. Here we provide evidence that CYFIP2 K/E editing has a role in neuronal axon and spine maturation, at least in primary neurons.

*CYFIP2* editing is mediated by ADAR2, and it is especially abundant in the cortex and cerebellar tissues [32]. A-to-I editing by ADARs also plays a role in embryogenesis and ageing, especially in the brain, highlighting its importance throughout the lifetime. In mice, RNA editing level of ADAR targets, including *CYFIP2*, increases through embryo development, and in some cases, it reaches almost 90% after 21 postnatal days [32,33]. Recently, it was shown that the levels of *CYFIP2* RNA editing are high only in neurons, together with a high level of protein expression [35]. *CYFIP2* RNA editing is undetectable in humans embryonic stem cells and foetal brains, while its level increases in the adult brain [36]. Interestingly, an age dependent decline in *CYFIP2* editing levels has been reported in the human adult brain [37]. Additionally, we have previously demonstrated that rat cortical cells treated with glutamate, the main excitatory neurotransmitter, show decreased *CYFIP2* RNA editing levels in parallel with a down-regulation of ADAR2 expression and self-editing [38]. Recently, modifications in *CYFIP2* editing levels have been linked to chronic social conflict [39]. Altogether, these reports show that *CYFIP2* RNA editing might be important for neural function and neuron maturation and further studies are needed to understand this relationship and the role of RNA editing in neurodevelopmental disorders.

The results reported here show that *CYFIP2* RNA editing reaction increases during neural development in a manner similar to other edited neuro-specific transcripts [18] and it is specifically regulated in different brain areas, suggesting a role during neuronal development and function. Furthermore, the level of CYFIP2 editing in hippocampal cultures varied under synaptic modulation suggesting that neuronal activity might functionally alter the proportion of CYFIP2 edited and unedited isoforms.

CYFIP2 protein is a fundamental component of Wave Regulatory Complex (WRC), a key hub for signalling between the plasma membrane and actin dynamic in a variety of functions. In its basal state, the WRC is inactive in the cytosol. After various upstream signals arising from growth factors, WRC can be recruited to the specific membrane regions where it activates the Arp2/3 complex to promote actin polymerization [38,39].

The interaction of Rac1 with the WRC is the primary driver of the complex activation. The active form of Rac (Rac-GTP) binds to the two sites of WRC (A and D) present in the CYFIP2 structure (or CYFIP1) and drives a series of conformational changes in WRC structure that leads to a release of the VCA domain which is free to activate the Arp2/3 complex to promote actin polymerization [42].

Accordingly, in our predicted 3D models, we observed that residue 320 is indeed located in an exposed portion of CYFIP2, pointing towards Rac1. Even if the measured distance is not sufficient to justify its involvement in a strong binding interaction, the variation of the residue from K to E leads to a small difference in terms of calculated binding energy, that anyway can be appreciated only if Rac1 is present, and that was not observed for the model of the inactive form.

To investigate if CYFIP2 K/E RNA editing regulation can be implicated in the control of actin dynamics, we initially used CYFIP2 KO neuroblastoma cell lines as an in vitro model. After CYFIP2 gene knockdown, SH-SY5Y cells lose their characteristic neuroblast-like phenotype, exhibiting a striking morphological change with the formation of cell blebs on the plasma membrane and a completely different actin filaments organisation compared to WT cells. To confirm the implication of CYFIP2 protein in actin cytoskeleton organisation, we create two stable SH-SY5Y CYFIP2 KI cell lines, one expressing the unedited CYFIP2 K variants one the edited CYFIP2 E variant. As expected, the alteration in morphological phenotype can be reverted when both CYFIP2 K/E variants are overexpressed. While these data clearly highlight the role of CYFIP2 in actin dynamics, however they do not clearly indicate a distinct effect of the edited variants.

Since CYFIP2 protein is mainly expressed in neurons [9] and the K/E RNA editing reaction is active only in the central nervous system, we investigate the role of CYFIP2 K/E variants on neuronal development. For this purpose, we use a model of neuronal differentiation based on the application on SH-SY5Y cells of retinoic acid and brain-derived neurotrophic factor [15] to monitor the conversion of undifferentiated cells into neuron-like cells. Differentiated WT cells assume a characteristic neuron-like morphology but in contrast, CYFIP2 KO cells display a distinctive fibroblast-like morphology with a non-polarized appearance and large soma. This evidence, suggesting the involvement of CYFIP2 protein in the process of neuronal development, is confirmed when we differentiate the CYFIP2 KI cell lines. We observe a complete restoration in the ability of neurite development in both CYFIP2 KI cell populations (K/E). However, a clear distinct function for the edited isoforms could not be described.

CYFIP2 has been shown to be involved in the growth and sorting of retinal ganglion cells axons [43]. For this reason, we initially focused on the early stages of neuronal maturation, using mouse hippocampal neurons, analysing axon length and complexity during the first days of in vitro neuronal development. Hippocampal neurons were co-transduced with lentiviral particles expressing CYFIP2 K or E variants together with lentivirus expressing shRNA to downregulate endogenous CYFIP2 protein, to assess the impact of RNA editing alteration, excluding the effect of endogenous protein. Increasing the expression of unedited CYFIP2 variant (K), leads to a simplification of axon development, in terms of the number of branching and axonal length, which is quite similar to the KD of CYFIP2. Conversely, an increase in the expression of the edited variant (E) results in a rise in axon complexity, even in comparison to WT neurons. Furthermore, we measured the frequency of the spines in secondary dendrites of mature neurons. The silencing of the CYFIP2 protein shows a decrease in the spine frequency compared to WT cells. This decrease remains similar when we induce in the expression of the K variant the cells, indicating that the physiological frequency of spines is not recovered after the expression of the CYFIP2 K variant. Conversely, this parameter was completely restored after the expression of the CYFIP2 E variant. Taken together these results suggest a clear role of CYFIP2 K/E RNA editing process in regulating both the outgrowth of neuronal axon during first stages of in vitro development and the process of spinogenesis in the in the subsequent stages of development of in vitro hippocampal cells. In both processes, the branching of b-actin fibers seems to be essential. Our findings suggest that the edited (E) variant of the CYFIP2 protein, when present within the WRC, might confer a gain of function in its ability to activate the WRC. Epi-transcriptomic A-to-I RNA editing can be temporally modulated to accurately modify the functions of neuronal genes throughout brain development. Thus, mediating numerous layers of controls necessary to build a sophisticated organ system like the brain. An increasing amount of evidence supports this idea, showing that RNA editing plays a crucial role in the regulation of central nervous system physiology [44]. During the process of neural differentiation and maturation and in all stages of brain development, from the embryo to the development of adult tissues, the levels of RNA editing of important neuro-specific transcripts dynamically change [43,44]. Our research has revealed for the first time how actin dynamic processes are related to the CYFIP2 K/E RNA editing process in the neuronal development and function. Further studies are necessary to determine whether this process is equally relevant in vivo.

## Supporting information

Supplemental results

## Acknowledgments

The authors declare that no funds, grants, or other support were received during the preparation of this manuscript. The authors performed experiments at the Imaging Platform of the Department of Molecular and Translational Medicine at the university of Brescia.

## Author Contributions

L.L.V. and A.B contributed to the study conception and design; L.L.V., E.N, M.B., V.M., G.C., A.F, F.B., G.R. acquired data; L.L.V., G.B., A.G., I.R., A.B. analyzed and interpreted the data; L.L.V. and A.B. prepared the draft of the paper; I.R, G.B. and C.F. critically revised the article. All authors approved final version of the article.

## Ethics approval

This study was performed in line with the principles of the Declaration of Helsinki. Approval was granted by the Ethics Committee of University of Brescia.

