## Supplemental results for "Functional Impact of CYFIP2 RNA Editing on Actin Regulation, Axon Growth, and Spinogenesis"

**Supplemental Figures**

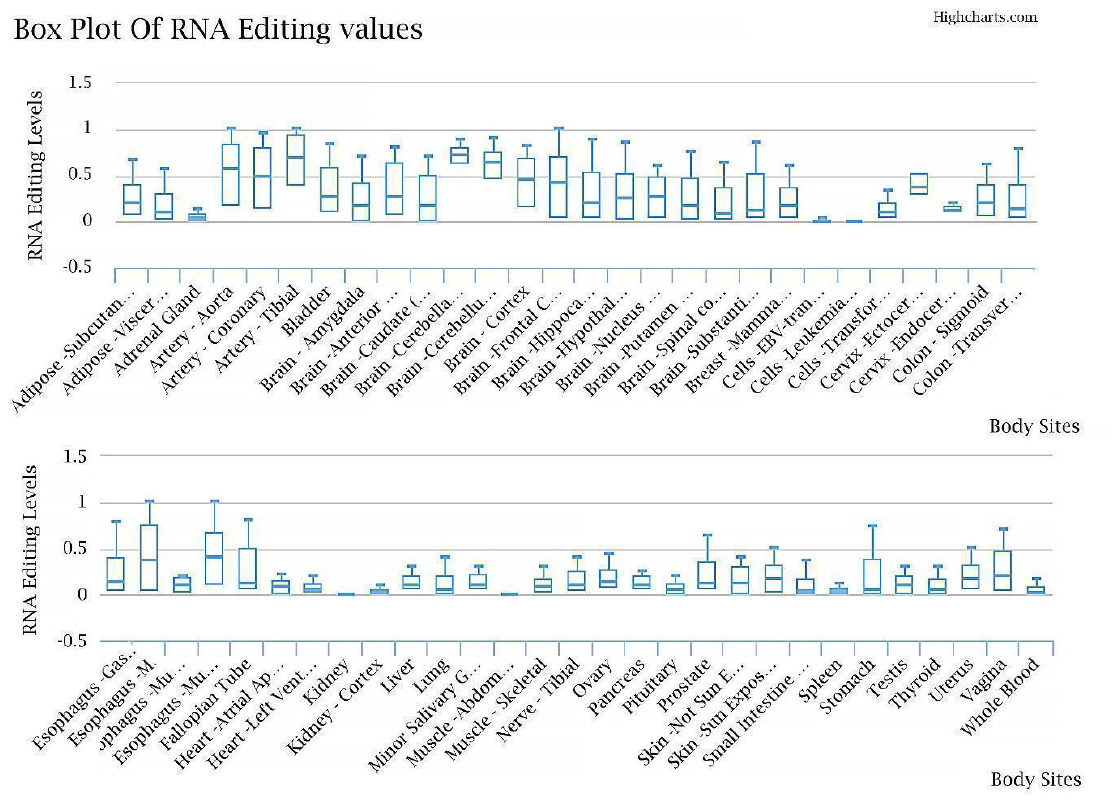

**Supplemental Figure 1**: RNA editing level of CYFIP2 K/E site in different human tissues as reported from REDIportal database (<http://srv00.recas.ba.infn.it/atlas/>).

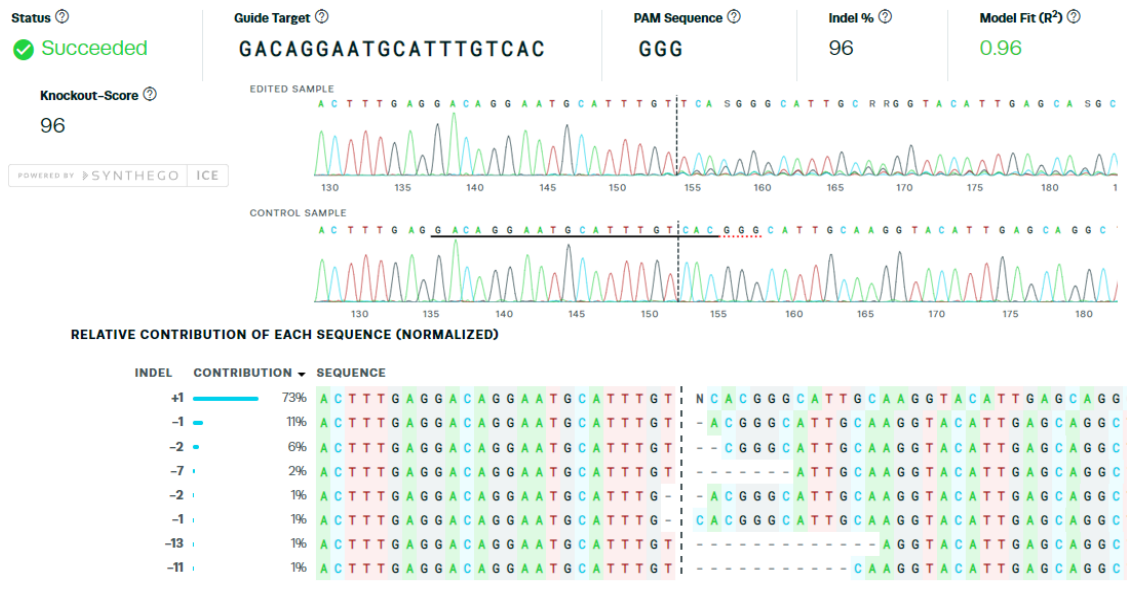

**Supplemental Figure 2**: Output obtained from Syntego© ICE Analysis tool. Comparison of electropherograms of the sgRNA recognition sequence (black underlined), obtained from sequencing of edited and WT samples (top). Alignment of different sequences present in the edited sample with relative contribution in percentage (bottom).

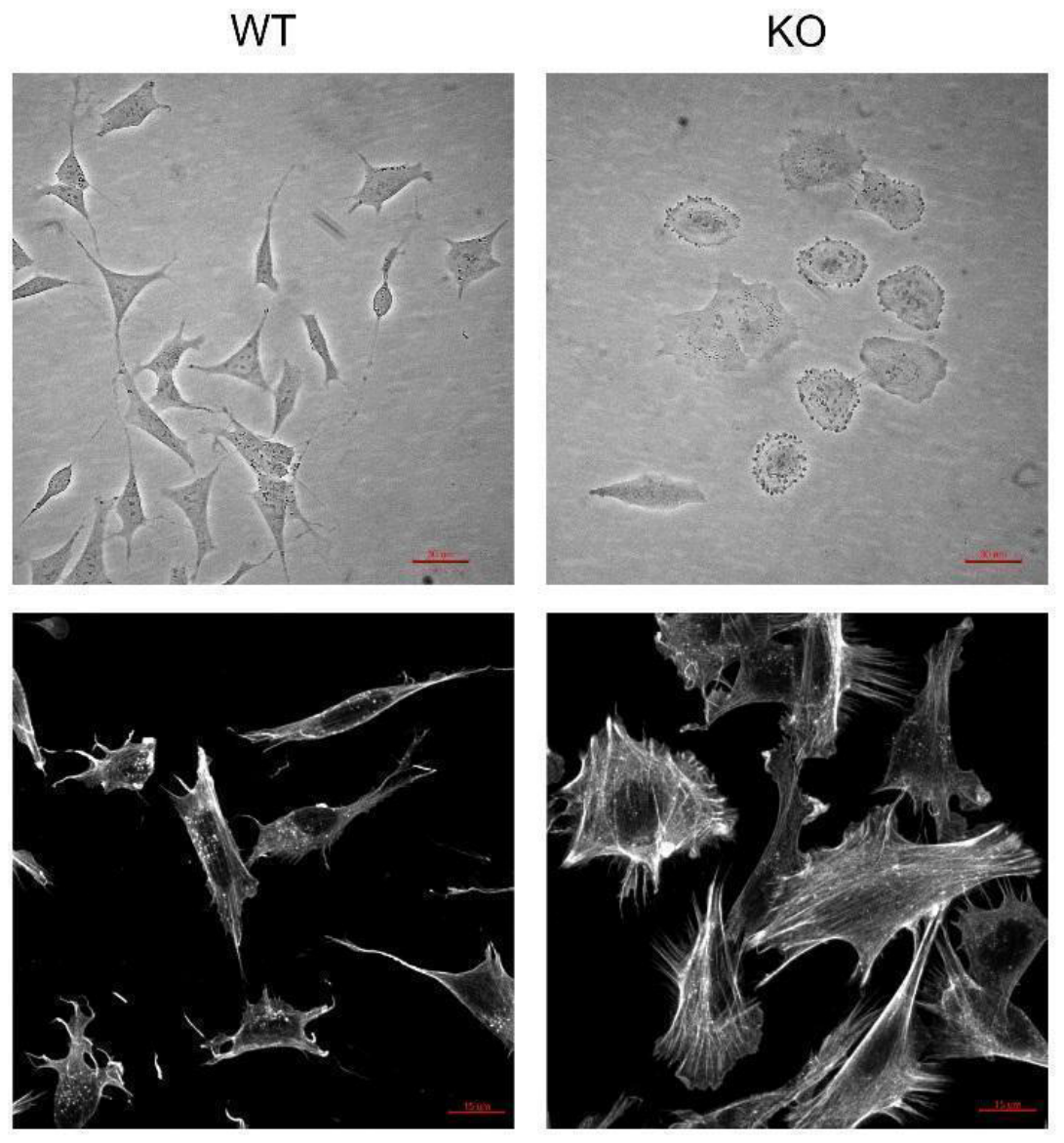

**Supplemental Figure 3**: Phase contrast microscopy images of SH-SY5Y wt (top, left) and SH-SY5Y KO (top, right) (Optech Biostar IB 20X scale bar: 30 μm). Phalloidin staining of SH-SY5Y cell line wt (bottom, left) and CYFIP2-KO (bottom, right) (Zeiss© LSM880 40X scale bar: 15 μm).
